# Weakly electric fish use self-generated motion to discriminate object shape

**DOI:** 10.1101/2022.12.01.518762

**Authors:** Sarah Skeels, Gerhard von der Emde, Theresa Burt de Perera

**Author notes:** Corresponding author (Sarah Skeels), Work address: The John Krebs Field Station, Department of Biology, University of Oxford, Oxford, OX2 8QJ, United Kingdom, Email address. Email addresses of other authors: Gerhard von der Emde, Theresa Burt de Perera. **Declarations of interest:** none. **Author contributions** Sarah Skeels: Conceptualisation, Methodology, Investigation, Formal analysis, Resources, Data Curation, Writing – Original Draft, Writing – Review & Editing, Visualisation, Funding acquisition Gerhard von der Emde: Conceptualisation, Methodology, Resources, Writing - Review & Editing, Supervision Theresa Burt de Perera: Conceptualisation, Methodology, Resources, Writing - Review & Editing, Supervision, Funding acquisition.

## Abstract

Body movements are known to play an active role in sensing. However, it is not fully understood what information is provided by these movements. The Peter’s elephantnose fish, *Gnathonemus petersii* sense their environment through active electrolocation during which they use epidermal electroreceptors to perceive object-induced distortions of a self-produced electric field. The analysis of electric images projected on their skin enables them to discriminate between three-dimensional objects. While we know the electric image parameters used to encode numerous object properties, we don’t understand how these images encode object shape. We hypothesise that ‘movement-induced modulations’ (MIMs) evoked by body movements might be involved in shape discrimination during active electrolocation. To test this, we trained fish to complete a shape discrimination task in a two-alternative forced-choice setup, and then manipulated the space available to individuals for scanning movements to see if this led to a change in their discrimination performance. We found that if enough space was available, fish were very good at discriminating objects of different shapes. However, performance decreased strongly when the space was reduced so that scanning movements were impaired. Our study demonstrates the importance of body movements for gaining complex environmental information such as object shape through active electrolocation. Movement can enhance perception by allowing the extraction of certain kinds of information. Similar observations have been made in other animals using different senses, suggesting that the core principles of sensory-motor integration might be valid for various sensory modalities.

---

Weakly electric fish, such as *Gnathonemus petersii (G. petersii)*, possess an active electric sense that they use to detect objects in their environment, termed active electrolocation (Lissmann, 1951, 1958; Lissmann & Machin, 1958; Heiligenberg, 1973). These fish generate electric organ discharges (EODs) from their electric organ. Each EOD produces a three-dimensional electric field around the fish. Objects within this field close to the fish’s skin (on the centimetre range) will distort it, and specialised electroreceptors on the skin will detect these changes (reviewed in von der Emde, 2006; von der Emde & Zeymer, 2020). The area of change on the skin is referred to as ‘the electric image’ of the object (Caputi & Budelli, 2006; von der Emde, 2006).

The active electric sense is well-suited to object recognition, as it provides fine scale spatial information of objects less than about half a fish length away (von der Emde e*t al*., 2010; Schumacher *et al*., 2016a, 2017a, 2017b). Individuals can determine object properties by examining different image parameters within an electric image, such as amplitude distribution within the image, local waveform distortion, image size, and others (von der Emde, 2006; von der Emde & Fetz, 2007; von der Emde & Zeymer, 2020). Thus, by analysing single electric images (‘snapshots’), it is possible for the fish to get information about an object’s size, material composition, distance, and electric colour (von der Emde *et al*., 1998; von der Emde & Fetz, 2007; Gottwald *et al*., 2017, 2018).

However, no parameter or collection of parameters in a single electric image can explain three-dimensional object shape (von der Emde, 2004, 2006; Schumacher *et al*., 2016b). Shape is a complex cue to extract from the environment, since it is made up of a collection of features that all contribute towards an object’s overall shape (e.g. object height, presence or absence of corners, the number, length, and orientation of sides; von der Emde & Fetz, 2007). Moreover, the perception of shape during electrolocation is, like in other senses, view-point dependent (Schumacher *et al*., 2016b; Fujita & Kashimori, 2019). In addition, the shape of an object at a given viewpoint cannot be taken as the shape of the corresponding electric image. There is no such mechanism within the electrosensory system to focus the electric image and the asymmetric nature of the electric field generated around the fish means that the electric image generated will always be distorted (Caputi & Budelli, 2006; von der Emde, 2006; Engelmann *et al*., 2008; Pusch *et al*., 2008). As such, there is no clear geometric relationship between the actual shape of an object and single electric images (Caputi & Budelli, 2006; von der Emde, 2006).

Nevertheless, we know that *G. petersii* can recognise three-dimensional shapes and accurately discriminate between them (von der Emde & Schwarz 2000, 2002; von der Emde, 2004; von der Emde & Fetz, 2007; von der Emde *et al*., 2010; Schumacher *et al*., 2016a, 2016b, 2017a; Skeels, 2022). They seem to use features within an object to recognise and generalise its shape, even when the object is rotated (von der Emde & Fetz, 2007; von der Emde *et al*., 2010; Schumacher *et al*., 2017a; Skeels *et al*., in prep). In the wild, it would be advantageous for *G. petersii* to determine an object’s shape accurately, so they can identify the object and determine the most appropriate response to it, e.g. an escape response away from a predator or an orientating movement during landmark detection.

Instead of using static electric images to determine the shape of an object, *G. petersii* might use the temporal series of electric images (electric flow) generated by engaging in movements around the object (Caputi & Budelli, 2006; von der Emde, 2006; Hofmann *et al*., 2013b). These movements will change the spatiotemporal dynamics of the electric field, and by analysing the changes in the electric field over successive electric images, they might be able to extract the necessary information required for shape recognition (Schumacher *et al*., 2016b; Fujita & Kashimori, 2019). We already know this kind of information can provide these fish with a cue for continuous distance estimation (Hofmann *et al*., 2017). Schumacher *et al*. (2016b) proposed that movement-induced modulations (MIMs) might act a cue for shape. MIMs refers to the ‘[temporal] changes in the electrical images that occur as a fish swims past an object’ (Schumacher *et al*., 2016b). They theorised that the magnitude and nature of these modulations would be dependent on the object’s shape and could provide a suitable cue for three-dimensional shape detection (Schumacher *et al*.,2016b). Although Schumacher *et al*. (2016b) found evidence to suggest a possible role of MIMs in shape recognition and discrimination, their limited sample size and inability to discount non-movement cues prevented them from drawing firm conclusions.

Our study aimed to test the role of MIMs in shape recognition by manipulating the motor component of fish behaviour. To do this, we trained fish in a two-alternative forced-choice (2AFC) setup to discriminate between two objects differing only in shape. Once the fish learned the task, we varied the space available for them to undertake scanning movements next to the objects -assuming that this negatively altered the amount of MIMs occurring- and tested whether this led to a change in discrimination performance. If MIMs were involved in shape discrimination, we would expect an individual’s discrimination performance to decline as swimming space –and thus MIMs-were restricted. Our study is therefore essential for determining the significance of body movements for three-dimensional shape recognition during active electrolocation in these fish.

## METHODS

### Subjects

We trained six *G. petersii* of unknown sex and age to complete a shape discrimination task in a 2AFC setup. We sourced our fish from a licenced fish dealer (The Goldfish Bowl, Oxford, UK). They had a total length of ca. 13.1-16.2 cm (including their chin appendage-Schnauzenorgan). Each fish was housed in a standard two-foot tank (approx. l × w × h: 60.2 × 35.2 × 31.5 cm) which also served as their experimental space. Reverse osmosis water was treated with a remineralisation formula to ensure fish well-being and stabilisation of the aquarium systems (Tropic Marin, Tropical Marine Centre, UK). Water conductivity was 415 ± 35 μS/cm, water temperature was 25.5 ± 0.5 °C, and pH 7.3± 0.3. We set the lights to a 12:12h light-dark cycle. Experiments were conducted during the light portion (20-30 lux). Fish were fed bloodworms (Gamma, Tropical Marine Centre, UK). Fish were only fed during training trials when participating in the study. Outside experiments, fish were given bloodworms twice a day, six times a week. We also supplemented their diet with brine shrimp (twice weekly) and additional vitamins (once a week). We provided plenty of enrichment (e.g. caves, tunnels, and plastic ornaments) to keep them engaged.

### Ethical note

This study was approved by the Department of Biology’s ethics committee and conducted in accordance with the University of Oxford’s animal ethics guidelines. No specific licences were required as the work undertaken was non-invasive. We took steps to ensure the well-being of our animals (see above) and limited the number of individuals kept in line with guidance provided by ASAB (Animal Behaviour, 2018).

### Experimental setup

Each tank was made up of four different compartments: the waiting area (l × w: 14.0 × 35.2 cm) where the fish began trials, the sensing area (26.2 × 35.2 cm) where the fish decided which object to choose, and the two experimental areas, A and B (each being 18.0 × 17.1 cm) which contained either the positive or negative object (Figs. 1 and 2). Outside of experiments, the fish could access all areas of their tank, but these now contained normal enrichment rather than experimental apparatus.

**Fig. 1.**
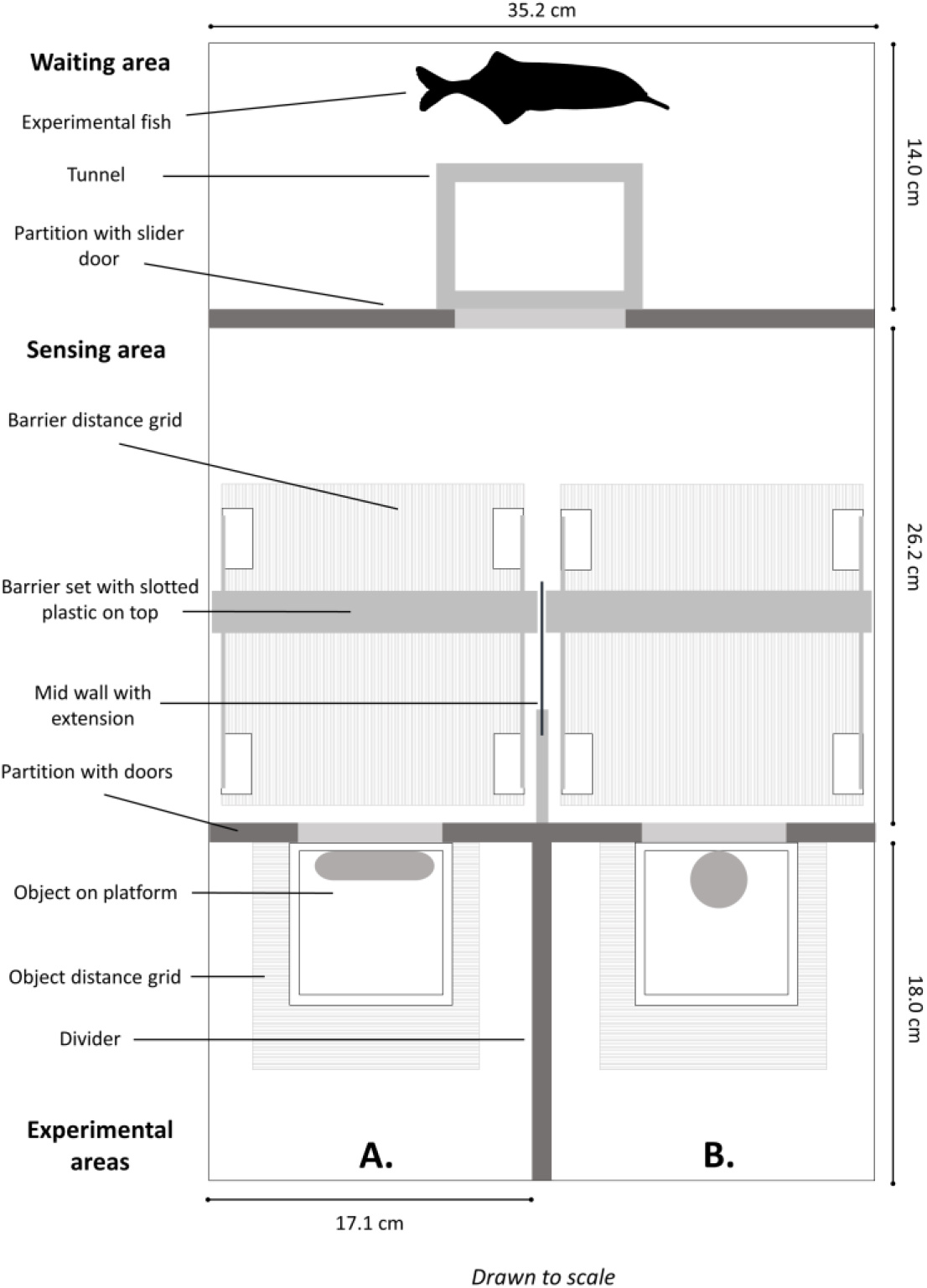
Top view of the experimental (2AFC) setup used during shape discrimination trials.

**Fig. 2.**
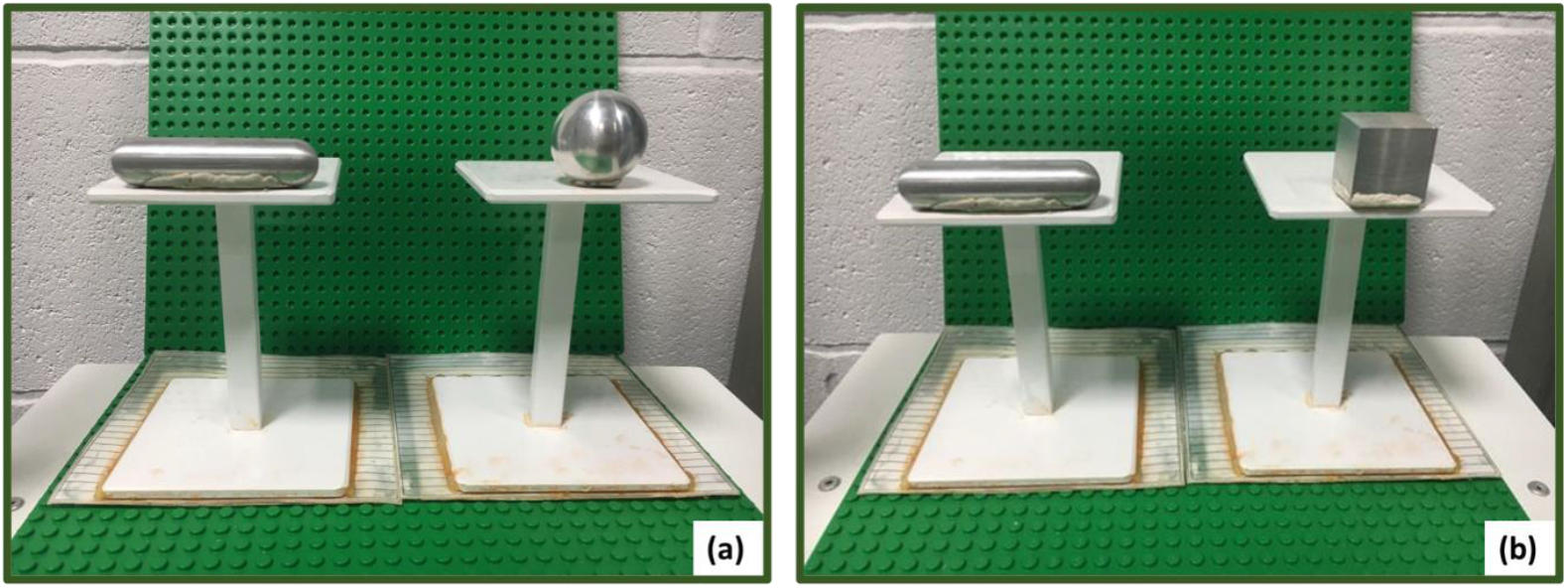
Objects used in the shape discrimination task. (a) Fish in group one (N=3) were trained to discriminate between a sausage-shaped object (S+) and sphere (S-). (b) Fish in group two (N=3) were trained to discriminate between a sausage-shaped object (S+) and a cube (S-). Objects only varied in their shape (material, volume, and distance were controlled for).

The partitions, barriers, and doors all had nylon mesh windows (mesh size: 0.71 mm per square) which meant that water could flow through them, ensuring that they did not form barriers for active electrolocation (Schumacher *et al*., 2016b). Barriers were placed in the sensing area to limit the amount of space that the fish could occupy and use for scanning movements during active electrolocation (Fig. 1). Two barriers were placed either side of the opening to A, and the same for B (Fig. 1). The overall size of each barrier was: h × w × d: 18.4 × 14.4 × 0.2 cm, and the mesh window was 14.8 × 7.9 cm. For training, the distance between each barrier pair was always 15.9 cm (D1), providing the maximal possible space. For testing, the distance was varied to one of four distances *(see testing regime)*.

### Pre-training

These sessions ensured the fish (N=6) were familiar with the setup before training fully commenced. They learned to swim out of the waiting area and into the sensing area when the slider door was opened. A clear tunnel (overall h × w × d: 28.9 × 10.9 × 7.7 cm) was placed at the entrance of this door to ensure the fish did not leave the waiting area at too much of an angle in case this had a bearing on choice (Fig. 1). They then learned to swim through the channels made by the barriers, open the doors leading to A and B, enter A/B to get a reward, and then return to the waiting area. They were also given time to acclimatise to the object platforms.

### Experimental Objects

The fish were split equally into two training groups. All fish were trained with the same positive object (S+) but each group was provided with a different negative object (S-). We did this to see if object pairing had an impact on performance. S+ was a sausage-shaped object (h × w × d: ca. 1.5 × 6.3 × 1.5 cm). S-was either a cube (side length 2.42 cm) or sphere (Ø 3 cm). The sausage-shaped object was presented with its longest side facing the door and the cube was presented with one of its sides facing the door (Fig. 2). All objects were made from the same material (aluminium) and of similar volumes, meaning that these properties should not have impacted choice (von der Emde & Fetz, 2007).

Each object was positioned on a platform (overall height ca. 10.8 cm). Objects were placed ca. 1.6 cm from the openings to A and B if fish were trained without the doors and ca. 1.9-2.5 cm away if trained with the doors (we had to account for door depth and water flow pushing doors out slightly here). Individuals moved the doors using their Schnauzenorgan. We trained and tested two individuals with the doors removed to check that the doors themselves did not influence performance (fish five and six).

### Training regime

They were trained to swim into the area with S+ (which was associated with a food reward-bloodworm) and to avoid the area with S-(which was associated with a mild punishment-a gentle tap of the glass followed by being shooed back to the waiting area). This combination of sound and movement was aversive to the fish but not enough to inflict stress.

The position of S+ was randomised to prevent choice being determined by object location. Generally, each fish completed 24 trials in a session. One session was conducted with each fish per day. A total of 3214 training trials were conducted across all fish during the training phase (fish one=312, fish two=480, fish three=264, fish four=655, fish five=629, and fish six=874 trials). An Apeman A70 Action Camera recorded the behaviour of the fish during all trials.

### Testing regime

Testing commenced once individuals had reached the pre-assigned learning criterion of 75% correct over three consecutive training sessions (as described in Schumacher *et al*., 2016a, 2016b; see appendix Fig. A1). Test trials were introduced every third trial. During these trials, the distance between the barriers was changed. The barriers could be placed at four possible positions: 15.9 (D1), 12.7 (D2), 9.5 (D3), or 6.3 cm apart (D4). Fish were neither rewarded nor punished to prevent further learning (von der Emde & Fetz, 2007; Schumacher *et al*., 2016b, 2017a). However, testing was interspliced with training trials (two for every test trial) to maintain discrimination performance and keep motivation high (von der Emde & Fetz, 2007). On average, we conducted 30 test trials at each distance for every individual studied. In total, we conducted 1460 training trials (fish one=240, fish two=240, fish three=240, fish four=240, fish five=248, and fish six=252 trials) and 720 test trials (fish one=118, fish two=118, fish three=120, fish four=117, fish five=123, and fish six=124 trials) in the testing phase alone.

### Posthoc analyses

#### Latency of response during test probes

Latency was defined as the time it took an individual to choose an object (measured from when the slider door was opened to when the fish had made their choice by entering either A or B completely). We were interested in looking at latency as a proxy for certainty. If individuals were using MIMs, we expected them to be more uncertain (and so take longer to make a choice) with the narrowing of the barriers, as scanning movements would be less effective in extracting shape information, and so more time inspecting the objects would be needed.

#### Side switching behaviour during test probes

Side switching was defined as the number of times an individual went between the two sides of the sensing area before making a choice. Side switching was examined as another proxy for certainty. If individuals were using MIMs, we expected them to be more uncertain (and so switch between the two sides more) when the barrier distance decreased, as scanning movements would be restricted, and so more inspections would be needed to discern the objects.

### Statistics

#### Shape discrimination performance

We used binomial generalised linear mixed models (GLMMs) to test for an effect of barrier distance on shape discrimination performance. Our main model contained two fixed effects (barrier distance and object pairing) and one random effect (fish ID). We investigated this model to determine what was driving the discrimination performance of our fish and the effect size of these predictors *(see appendix for further details)*.

#### Latency of response during test probes

We used gamma GLMMs to determine whether barrier distance had a significant effect on latency. Our main model had two fixed effects: barrier distance and trial outcome (which we hypothesise would be correlated with level of certainty), with the random effect set as fish identity. The model was investigated to determine what was influencing response times and the strength of the influence *(see appendix)*.

Both analyses were run in *R Studio, R version 3*.*6*.*2* using the ‘lme4’ package (Bates *et al*., 2015).

#### Side switching during test probes

We used Poisson GLMMs to determine whether barrier distance had a significant effect on side switching. Our main model stated that side switching depended on barrier distance and trial outcome (fixed effects) and fish identity (random effect). We examined the model to determine what was influencing side switching and the strength of this influence *(see appendix)*.

This analysis was run in *R Studio, R version 4*.*1*.*2* using the ‘lme4’ package (Bates *et al*., 2015).

## RESULTS

### Shape discrimination performance

We found that an individual’s ability to discriminate differently shaped objects was significantly influenced by how far apart the barriers in the sensing area were (GLMM: *Z*_716_=-6.184, *P*<0.001; Figs. 3 and A2; Tables A1, A2, and A3). All fish learned to discriminate between the two objects offered (Fig. A1). However, they generally found it harder to correctly identify the positive object as the distance between the barriers was reduced (Figs. 3 and A2; Table A1). This was regardless of which objects the fish were trained and tested with (GLMM: *Z*_716_=0.795, *P*=0.426). Door presence/absence did not seem to influence discrimination performance (Fig. A2).

**Fig. 3.**
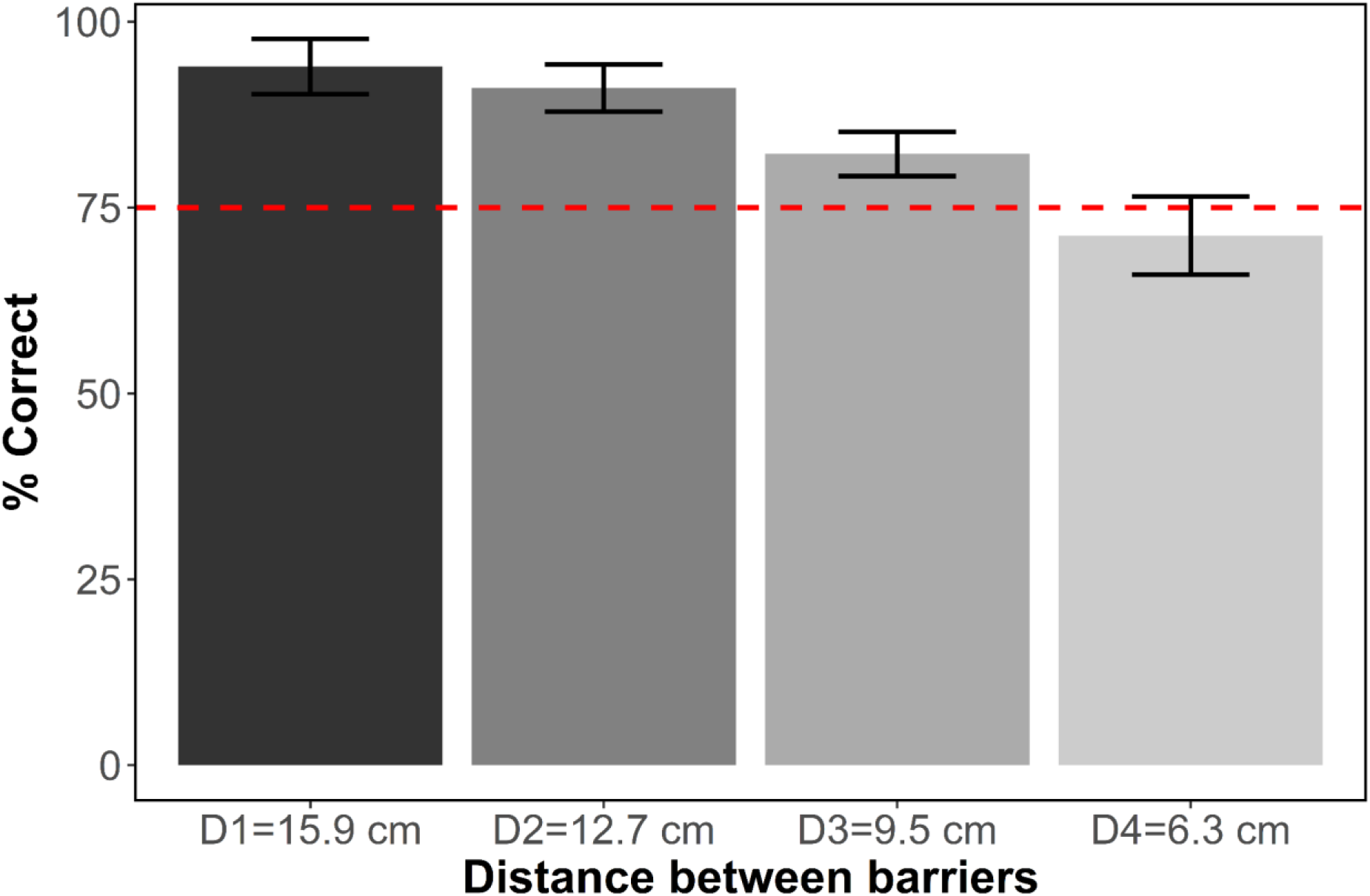
Group mean discrimination performance (with SE bars) during the testing phase. Performance was measured as the percentage of trials that were correct for a given distance (D1-D4). The dotted line denotes the learning threshold of 75% correct. Total N trials=720 (for an individual breakdown *see testing regime in methods)*.

### Posthoc analyses

#### Latency of response during test probes

We found that barrier distance was a significant predictor of latency (GLMM: *T*_703_=-19.949, *P*<0.001; Fig. 4). Latency increased as barrier distance decreased, with the most noticeable increasing occurring at the narrowest distance, D4 (mean latency ± SE: D1=6.88 ± 0.36 s, D2=6.74 ± 0.28 s, D3=9.25 ± 0.48 s, and D4=26.19 ± 2.60 s; Fig. 4). Trial outcome did not have a significant effect on latency (GLMM: *T*_703_=0.677, *P*=0.499; Fig. 4).

**Fig. 4.**
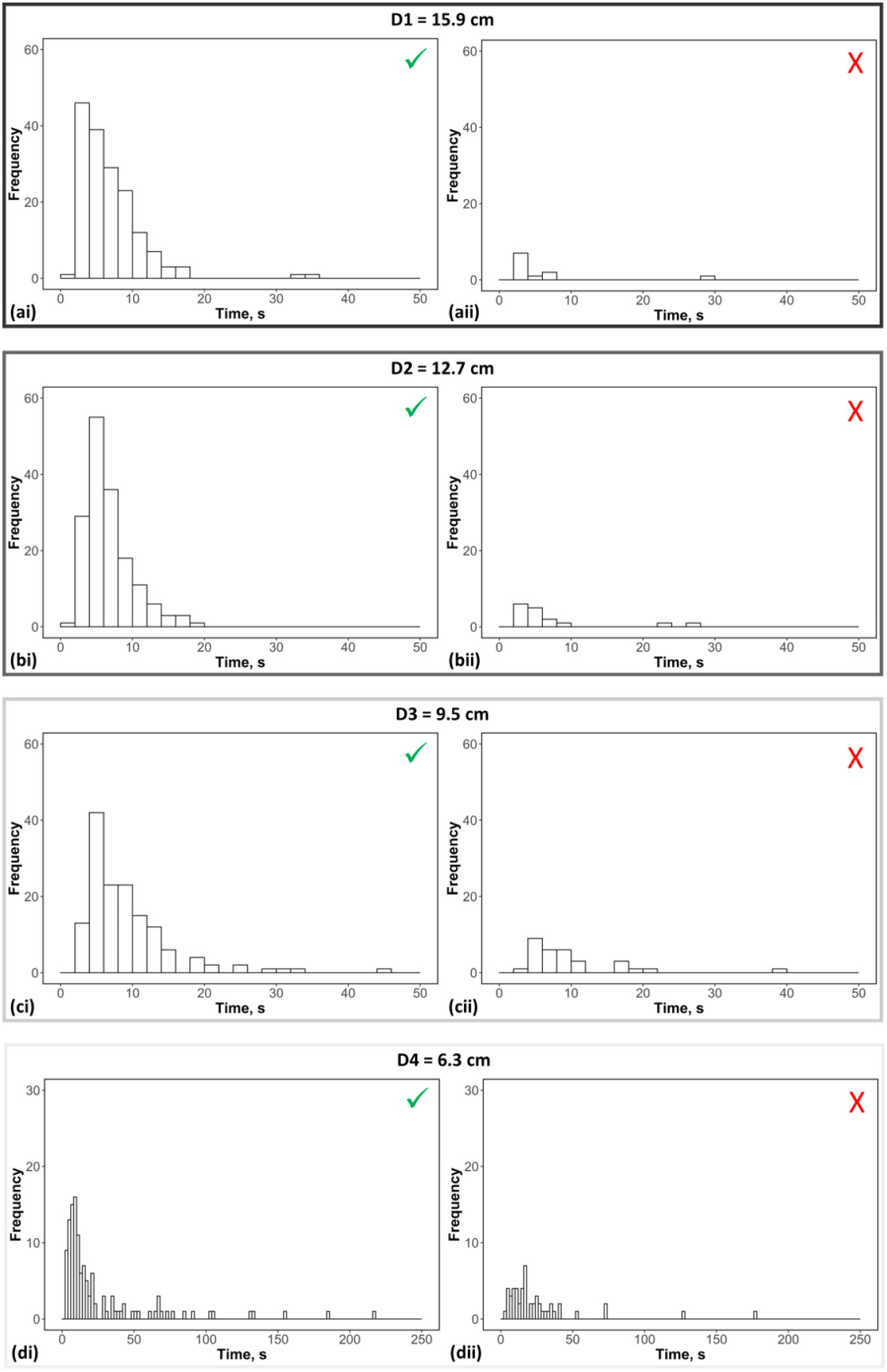
Response times for completing the shape discrimination task during test probe trials at D1-D4 (a-d). Each data point represents the time it took an individual to make a choice in a single trial (given in seconds, s). Times have been pooled to show responses on a group level. For each distance, response times have been categorised according to whether the trial resulted in a correct (i; tick) or incorrect choice being made (ii; cross). Note that the same x-axis range has been used for D1-D3, but a larger x-axis range was required for D4 due to the steep increase in response times. Bin size was the same for all conditions to aid comparisons. N=708 trials (ai=165, aii=11, bi=163, bii=16, ci=146, cii=31, di=125, and dii=51).

#### Side switching behaviour during test probes

Individuals tended to prefer to limit their side switching where possible, with most trials showing no side switching or only a single side switch (Figs. 5 and A3). Nevertheless, we found that barrier distance had a significant influence on side switching behaviour (GLMM: *Z*_716_=2.905, *P*=0.0037). More switching occurred as barrier distance was narrowed-particularly at the shortest distance, D4 (Fig. 5). The amount of switching did vary across individuals (Fig. A3). Fish five and six showed lower levels of multiple switching compared to the other fish studied, and both were trained and tested without the doors (Fig. A3). Lastly, trial outcome did not have a significant influence on side switching (GLMM: *Z*_716_=0.725, *P*=0.4686; Fig. 5).

**Fig. 5.**
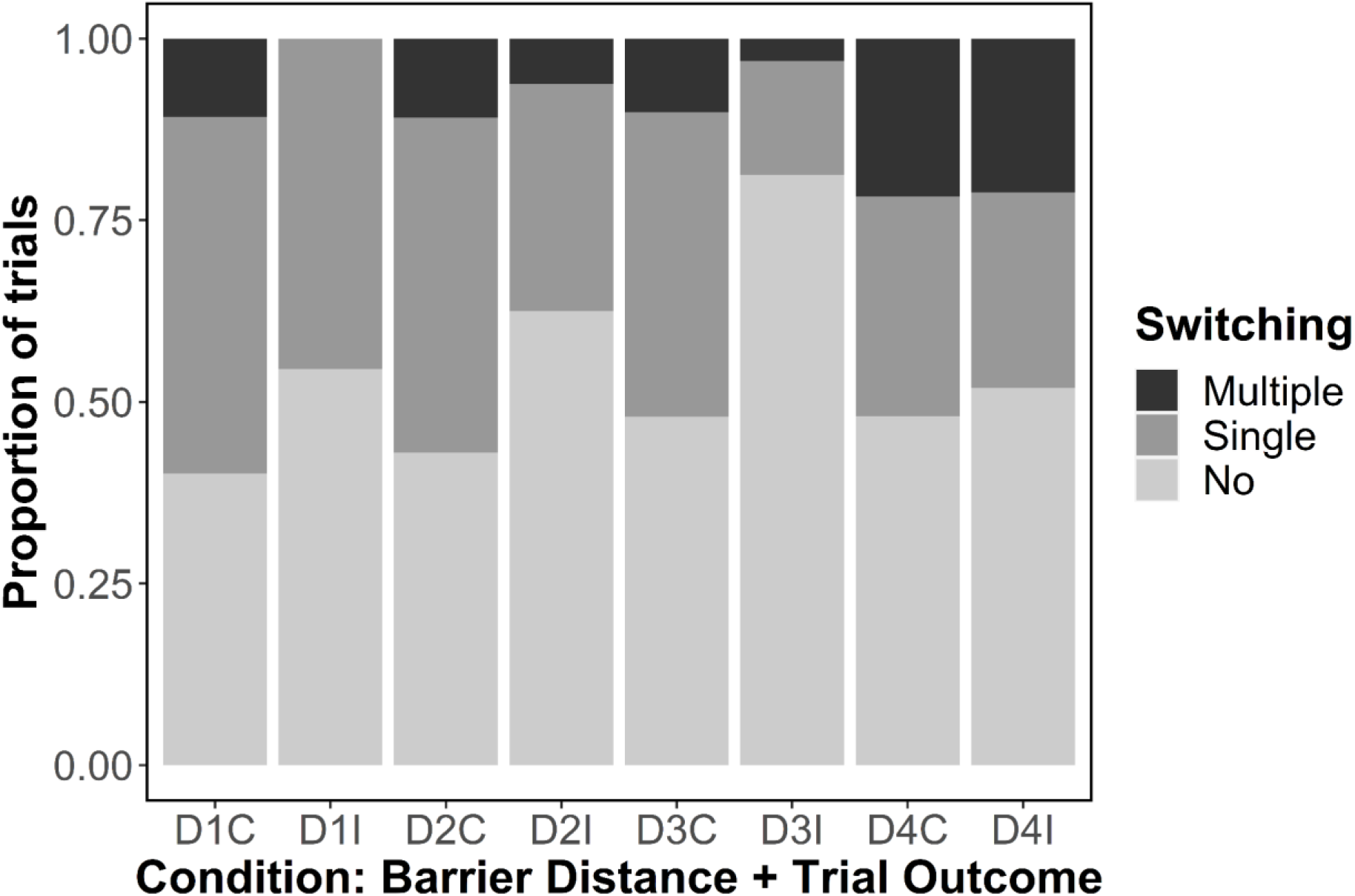
Group side switching behaviour during test probe trials. Trials were placed into one of eight conditions. The condition was dependent on the barrier distance used for that trial (D1-D4) and the outcome of the trial (C=correct trial and I=incorrect trial). Greyscale bars indicate the type of side switching behaviour observed (lightest grey=no side switching, middle grey=single side switch, and lastly, black=multiple (two or more) side switches). The size of each bar represents the proportion of trials which fitted each category. Total N trials=720 *(for an individual breakdown see appendix)*.

## DISCUSSION

We investigated the role of egocentric movement in shape discrimination in *G. petersii*. We were interested in determining whether movement-induced modulations (MIMs) could be a useful method for extracting shape information. To determine this, we trained fish to complete a shape discrimination task in a 2AFC setup. We manipulated the space available for scanning movements to determine whether this had an impact on their ability to distinguish the objects correctly.

### Shape discrimination performance

We found that barrier distance had a significant effect on an individual’s ability to discriminate between the two objects (Figs. 3 and A2; Table A1). Individuals found it more difficult to discriminate the objects as the barrier distance narrowed and space around the object became more limited-regardless of the objects they were trained and tested with (Figs. 3 and A2; Table A1). This supports our hypothesis that *G. petersii* use movement to facilitate the extraction of shape information from the environment.

We found that the biggest drops in discrimination performance tended to occur at the shortest barrier distances (Figs. 3 and A2; Table A1). These drops likely represent a threshold being reached, where the amount of space available was not enough to reliably scan the objects with the active electric sense, and so lacking information on the objects’ shape, mistakes became more frequent.

The hypothesis that the barriers shaped the movement of the fish is further supported by our observations at the shorter distances. We found individuals tended to change their movements when beside the objects, and this was seen across all fish, regardless of whether or not they were trained with the doors, suggesting the changes were not due to the presence or absence of doors but the barriers themselves *(see appendix-Notable behaviours during testing phase)*.

Given that we did not block out any senses due to the impact on behaviour, how can we be sure that our fish are primarily using their active electric sense for shape discrimination and not another sense? In principle, an individual could get shape information about the two objects using vision (e.g. Schumacher *et al*., 2016a). Since the two objects would have looked very different when observed from straight ahead, a reduction in space between the barriers should not have impaired visual discrimination. Movement in front of the objects would not have provided additional information and thus not helped visual discrimination. For electrical discrimination, however, movement might have been necessary. Since the objects had the same volume, they would have elicited quite similar electric images when a fish swam straight ahead towards them without left and right movement (Fujita & Kashimori, 2019). Only when the fish was moving in front of the objects would the electric image have changed significantly, according to the angle between the fish and the object and would have also moved on the skin of the fish until the objects had been electrolocated from the sides (Hofmann *et al*., 2013a, 2013b). These temporal changes of electric images and their movement on the skin of the fish might have been necessary to get enough information about an object’s shape (Hofmann *et al*., 2013a, 2013b; Schumacher *et al*., 2016b; Fujita & Kashimori, 2019). So, if we impair movement of an individual, they might only get straight ahead electric images, which do not change temporally and do not move on the skin (or move less), and therefore may not provide enough information to discriminate between the two objects. Given our findings that discrimination performance declined with barrier distance, and the dominance of the active electric sense for object recognition at the distances we tested at (e.g. Schumacher *et al*., 2017a), we can be confident that these fish were primarily relying on their active electric sense when completing our shape recognition task.

In summary, we found that shape discrimination performance worsened as barrier distance declined, suggesting that individuals extracted shape information through movement, presumably from analysing successive electric images as the fish moved alongside the object, i.e. via MIMs (von der Emde *et al*., 2010; Fechler & von der Emde, 2013; Hofmann *et al*., 2013b; Schumacher *et al*., 2016b; Fujita & Kashimori, 2019).

### Posthoc analyses

#### Latency of response during test probes

We examined response times as an indicator of the level of certainty an individual experienced during trials under a set number of conditions. Latency increased as barrier distance decreased (Fig. 4), suggesting that individuals were more uncertain during trials at shorter distances, and so needed more time examining the objects before they could make a choice. This is likely because a discrimination threshold with MIMs was reached. This would have occurred when the space was restricted to such a degree that an individual was unable to scan the objects effectively with their active electric sense, meaning that extra scans were required to resolve any ambiguities that existed between the two objects. In other words, when the barriers were at their most restrictive, the fish were less successful at scanning the objects, so they tried harder (see below) and for a longer time.

#### Side switching behaviour during test probes

We measured switching behaviours as another proxy of certainty. We expected individuals to perform fewer side switches (and take less time) when they were more certain of S+ but exhibit more side switches (and take more time) when they were less certain. Fish completed more side switches when the distance between the barriers narrowed (Fig. 5), supporting the hypothesis that the fish were primarily relying on MIMs to complete the shape recognition task, and that side switching was employed to add value in trials, for example to help resolve ambiguities between the two objects in situations of greater uncertainty through additional scans. This is supported by von der Emde and Fetz (2007) who found that individuals who inspected both objects had a greater amount of success in picking S+ than those that examined just one object. As such, side switching likely played an important role in our study by minimising the chance of the wrong object being chosen.

Side switching may also have been a consequence of individuals resorting to using other cues (e.g. vision) if MIMs were compromised. Fish using these other cues would have likely needed more time and visits with the objects before having enough information to make an informed choice, and even that would not have guaranteed success, as these cues would not have been optimised for fine scale object recognition-like vision (Landsberger *et al*., 2008; Kreysing *et al*., 2012; Pusch *et al*., 2013a, 2013b; Francke *et al*., 2014).

#### Conclusions

Here, we show that movement is key to enabling *G. petersii* to recognise and successfully discriminate between objects differing in their shape. We found that performance worsened as the barrier distance was decreased. Our analysis supports the hypothesis that our fish were predominately relying on their active electric sense for shape discrimination, and that movement-induced modulations (MIMs) of a series of electric images moving over the fish’s skin generated by swimming was likely the cue being used. Our study therefore contributes to the growing body of evidence that movement is essential for shaping sensory input in a wide range of taxa (see reviews: Land, 1999; Caputi, 2004; Nelson & MacIver, 2006; Mitchinson *et al*., 2011; Prescott *et al*., 2011; Hofmann *et al*., 2013b; Yang *et al*., 2016; Engelmann *et al*., 2021). Animals use movement in similar ways to enhance their perception of the environment regardless of their taxonomic group (e.g. Prescott *et al*., 2011) or preferred sensory modality (e.g. Hofmann *et al*., 2013b). This indicates the fundamental importance of sensory-motor coupling to understanding animal behaviour generally. We suggest future studies might benefit from taking a comparative approach in order to interpret and understand these movements better.

## Acknowledgements

We would like to thank Oliver Padget for their statistical advice, Adélaïde Sibeaux for their helpful comments on this manuscript, as well as David Skeels and Toby Skeels-Jungius for proofreading. Christine Soper and Helen Sanders-Parker provided invaluable assistance with fish husbandry, and John Hogg built the 2AFC setups used. This work was supported by funding from the Biotechnology and Biological Sciences Research Council (BBSRC) [grant number BB/M011224/1]. Sarah Skeels is currently funded by a BBSRC Career Development Fellowship.

We would like to thank XXXX for their statistical advice, XXXX for their helpful comments on this manuscript, as well as XXXX and XXXX for proofreading. XXXX and XXXX provided invaluable assistance with fish husbandry, and XXXX built the 2AFC setups used. This work was supported by funding from the Biotechnology and Biological Sciences Research Council (BBSRC) [grant number BB/M011224/1]. XXXX is currently funded by a BBSRC Career Development Fellowship.

## Appendix

### GLMM information

#### Shape discrimination performance during testing

We used binomial generalised mixed models (GLMMs) to test for an effect of barrier distance on shape discrimination performance. We initially created two models to predict correct object selection, the first contained two fixed effects (barrier distance and object pairing) and one random effect (fish ID). The second model dropped our main parameter of interest (barrier distance). We compared the models using a likelihood-ratio test (LR) to determine which one was a significantly better fit to our data. We found that our first model best fitted the data (LR test: *χ*^*2*^_1_ =43.403, *P*<0.001). We then investigated this model to determine how barrier distance influenced the discrimination performance of our fish and the effect size of this predictor.

#### Latency of response during test probes

We used gamma GLMMs to determine whether barrier distance had a significant effect on latency. We initially created two models. Our first model had two fixed effects: barrier distance (our key parameter) and trial outcome (we thought this be correlated with level of certainty). Our random effect was fish identity. The second model omitted barrier distance. A LR test was then done to compare the models to assess which one best fitted our data. We found that model one best described the latency data (LR test: *χ*^*2*^_1_ =378.79, *P*<0.001). This model was then run and investigated to determine what was influencing response times and the strength of the influence.

#### Side switching during test probes

We used Poisson GLMMs to determine whether barrier distance had a significant effect on side switching. We created two models, our first, stated that side switching depended on barrier distance and trial outcome (fixed effects) and fish identity (random effect). The second model omitted barrier distance as a predictor. A LR test compared the two models to assess which one was a better fit. We found that model one fit the data better (LR test: *χ*^*2*^_1_ =8.434, *P*=0.0037). We then examined this model to determine what was influencing side switching and the strength of this influence.

#### Random effects

Our random effect was fish ID for all analyses. For the performance model, our random effect was σ^2^=0.212 and σ=0.460. For the latency model, our random effect was σ^2^= 0.001 and σ=0.024, and for the side switching analysis, our random effect was σ^2^=0.115 and σ=0.340.

#### Notable behaviours during testing phase

During training and test probe trials, *G. petersii* displayed particular behaviours whilst investigating the objects behind doors A and B. When leaving the waiting area, individuals tended to head to one side first, often the side that they had a pre-existing preference for. This preference seemed to be specific to an individual and likely developed over time. During their approach, the head, Schnauzenorgan, and body were held out relatively straight and level in the water column. Once beside the objects, they adjusted their body and head positions frequently and were often more angled. We observed several types of ‘probing motor acts’ during trials (Toerring & Belbenoit, 1979; Toerring & Moller, 1984; von der Emde & Fetz, 2007). Head/stationary probing and chin probing were observed when the fish were close to the objects. Tail probing was observed when the fish were unfamiliar with an object or seemed wary of it. Tangential probing occurred when the fish examined one object before rapidly changing direction to explore the other object. Whole-body back-and-forth *(‘va-et-vient’)* motions were less observed than other motor acts due to the constraints on space. When barriers were at their most restrictive (D4), fish tended to inspect the objects from more directions and angles; there were lots of small adjustments to the head and body. They also adjusted their door opening technique. Fish trained without the doors also showed similar shifts in the way they used the space around the objects.

**Table A1.**
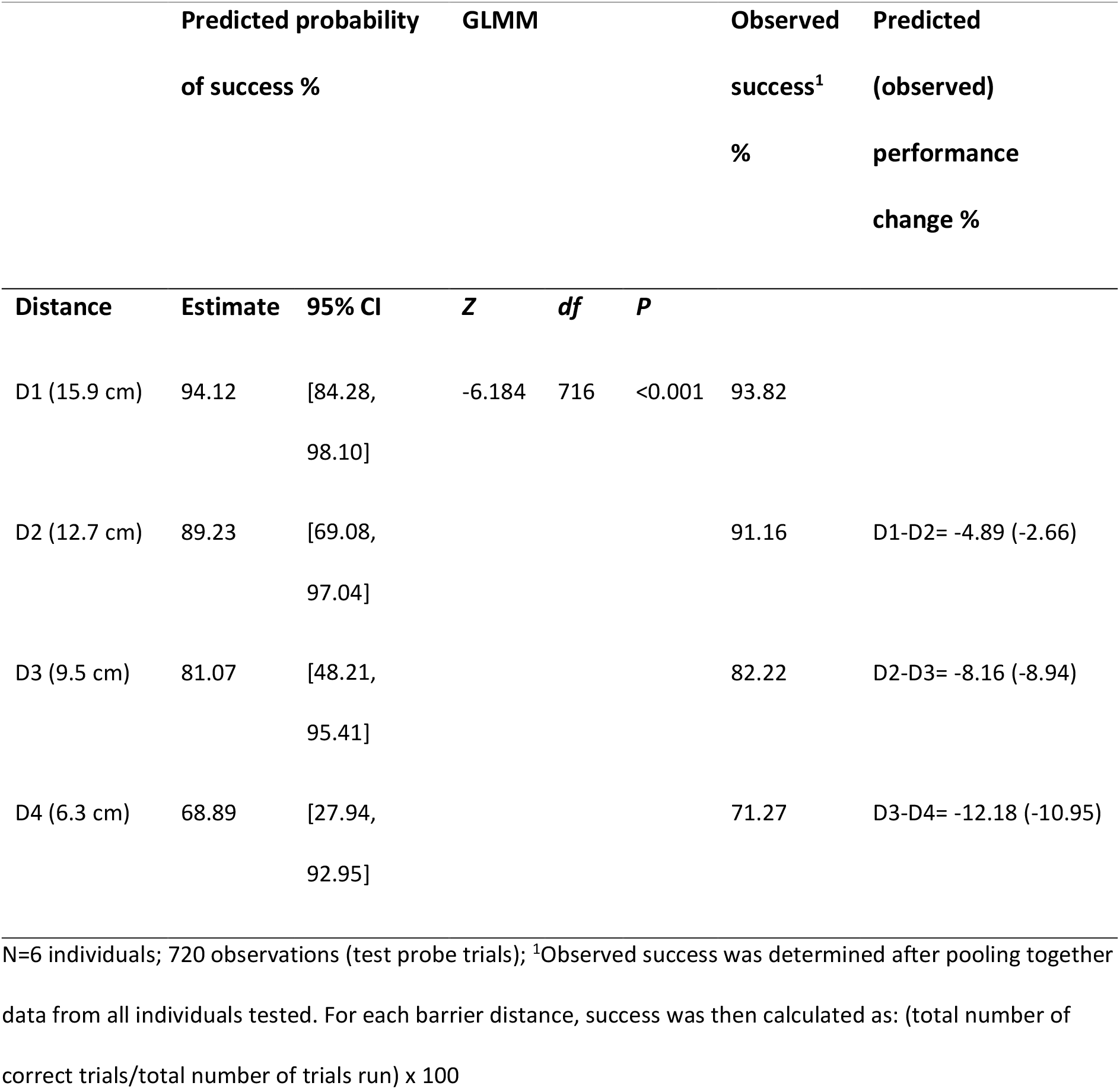
Predicted and observed values of success for a test trial given barrier distance

**Table A2.** Values for the estimate derived from the model that best described discrimination performance

| Distance | Log odds ratio | Odds ratio | % likely to get correct | Performance step change (%) |
| --- | --- | --- | --- | --- |
| D0(intercept) | 3.433 | 30.983 | 96.87 |  |
| D1: 15.9 cm | 2.774 | 16.020 | 94.12 | D0-D1: -2.75% |
| D2: 12.7 cm | 2.114 | 8.283 | 89.23 | D1-D2: -4.89% |
| D3: 9.5 cm | 1.455 | 4.283 | 81.07 | D2-D3: -8.16% |
| D4: 6.3 cm | 0.795 | 2.214 | 68.89 | D3-D4: -12.18% |

**Table A3.**
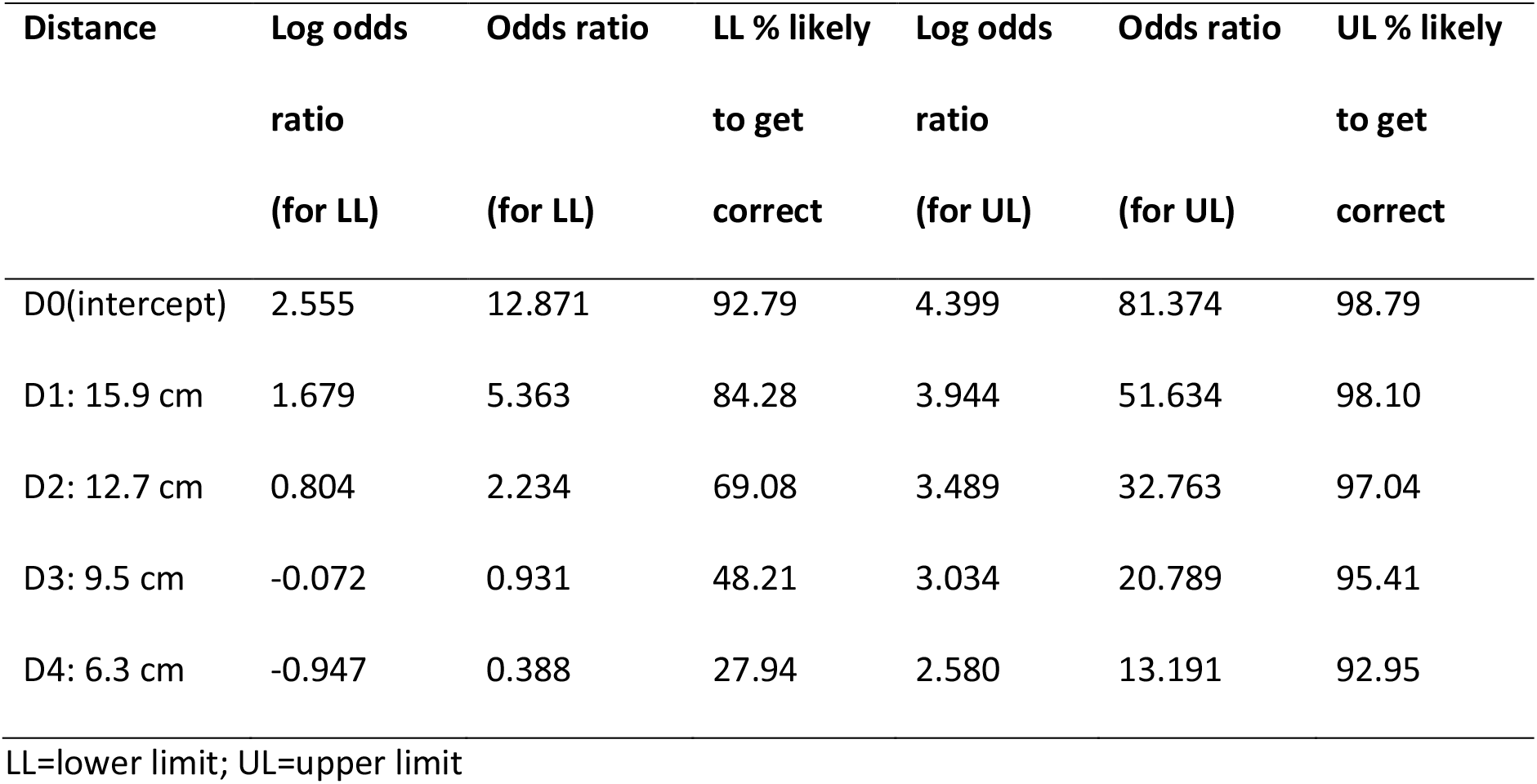
Values for the lower and upper limits derived from the model that best described discrimination performance

| Distance | Log odds ratio<br>(for LL) | Odds ratio<br>(for LL) | LL % likely<br>to get<br>correct | Log odds ratio<br>(for UL) | Odds ratio<br>(for UL) | UL % likely<br>to get<br>correct |
| --- | --- | --- | --- | --- | --- | --- |
| D0(intercept) | 2.555 | 12.871 | 92.79 | 4.399 | 81.374 | 98.79 |
| D1: 15.9 cm | 1.679 | 5.363 | 84.28 | 3.944 | 51.634 | 98.10 |
| D2: 12.7 cm | 0.804 | 2.234 | 69.08 | 3.489 | 32.763 | 97.04 |
| D3: 9.5 cm | -0.072 | 0.931 | 48.21 | 3.034 | 20.789 | 95.41 |
| D4: 6.3 cm | -0.947 | 0.388 | 27.94 | 2.580 | 13.191 | 92.95 |

LL=lower limit; UL=upper limit

**Fig. A1.**
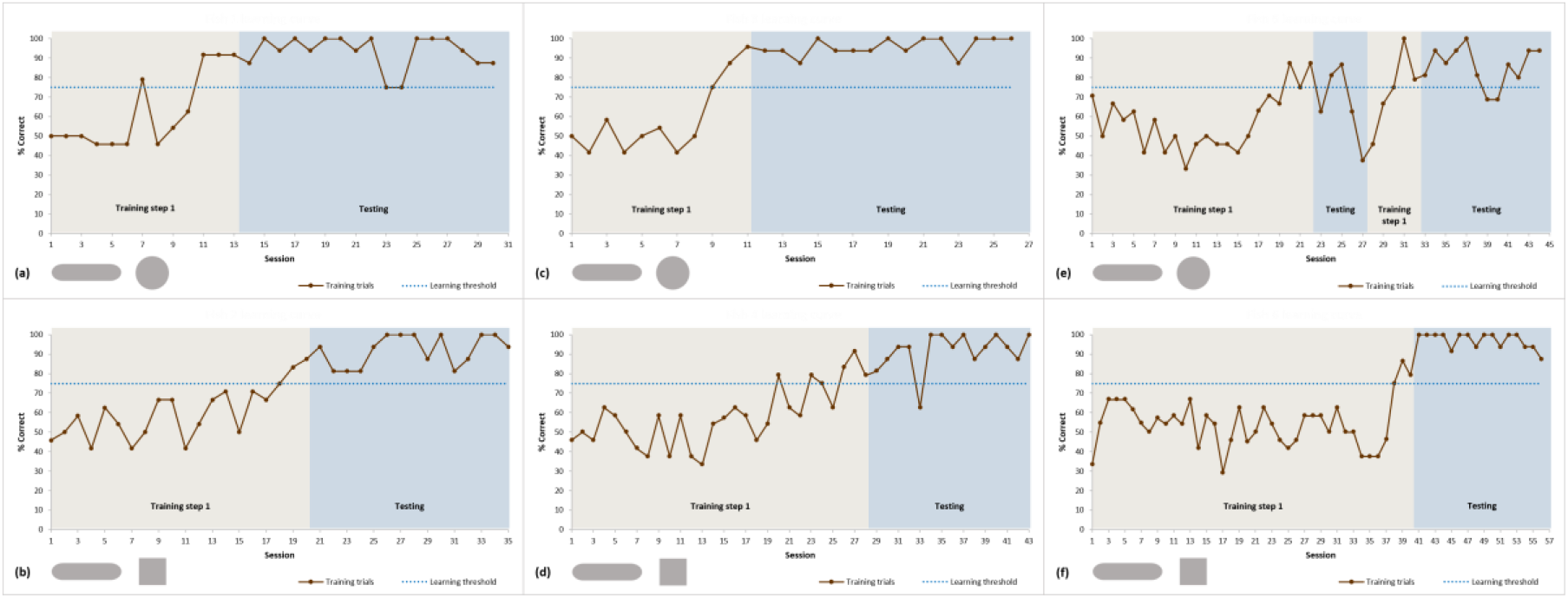
Learning curves for fish one to six (a-f). Fish were trained to discriminate between two differently-shaped objects. S+ (a sausage-shaped object) was the same for all fish. Fish one (a), three (c), and five (e) had the sphere as their S-, whereas fish two (b), four (d), and six (f) had the cube. Fish one to four were trained with doors, whereas fish five and six were not. During training step one, an individual was trained to distinguish between S+ and their S-(shaded in light brown). Once the fish had reached or exceeded the learning threshold of 75% on three consecutive sessions (dotted line) they moved onto testing (shaded blue). During testing, training trials were interspliced with test probes to maintain motivation and performance. The scores from these training trials are plotted here. Total N=4722 training trials across both experimental phases (fish one=584, fish two=720, fish three=504, fish four=895, fish five=893, and fish six=1126).

**Fig. A2.**
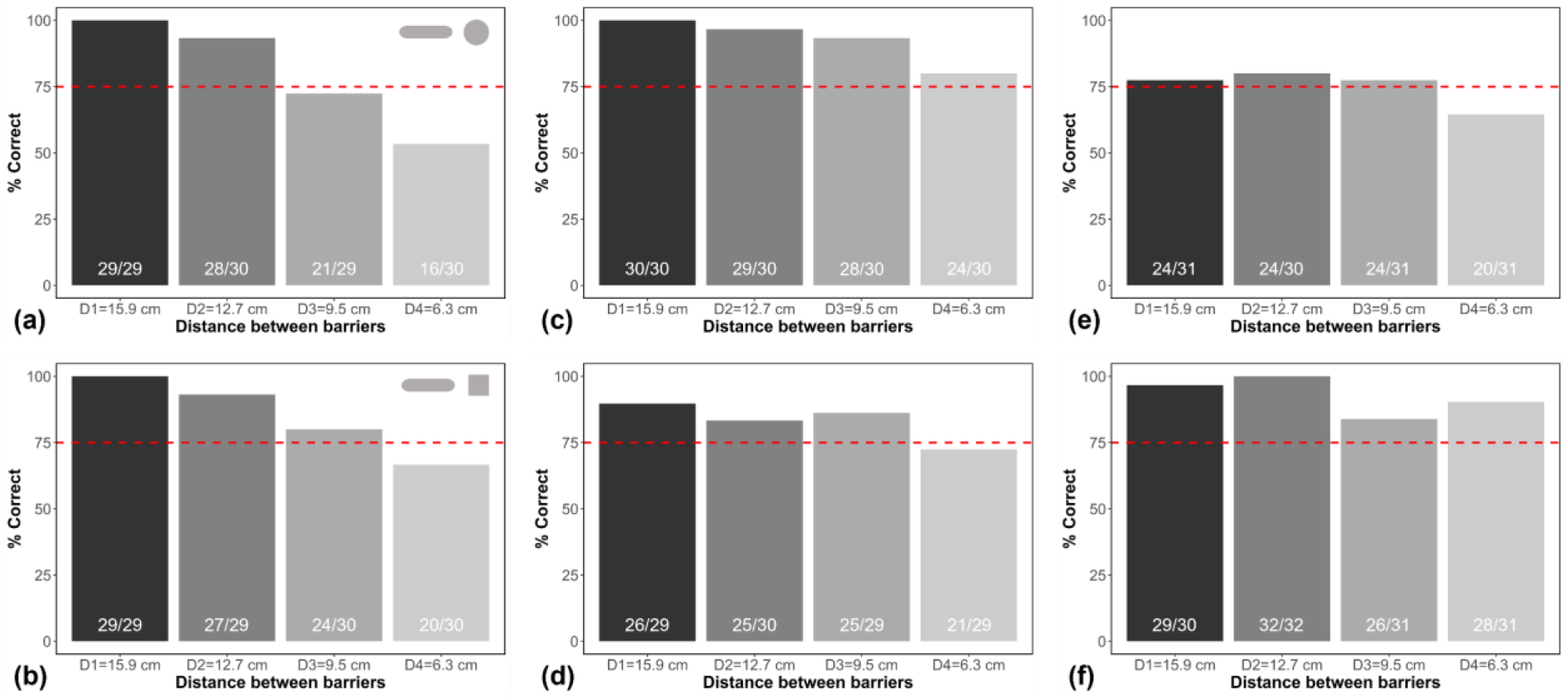
Individual discrimination performance during the testing phase (a-f). Fish one, three, and five were all trained with the sphere as their S-(a, c, and e). Fish two, four, and six had the cube as their S-(b, d, and f). The numbers within each bar show how many trials were correct against how many were run. The dotted line represents the learning threshold of 75%. Note that fish five and six (e and f) were trained and tested without the doors by A and B.

**Fig. A3.**
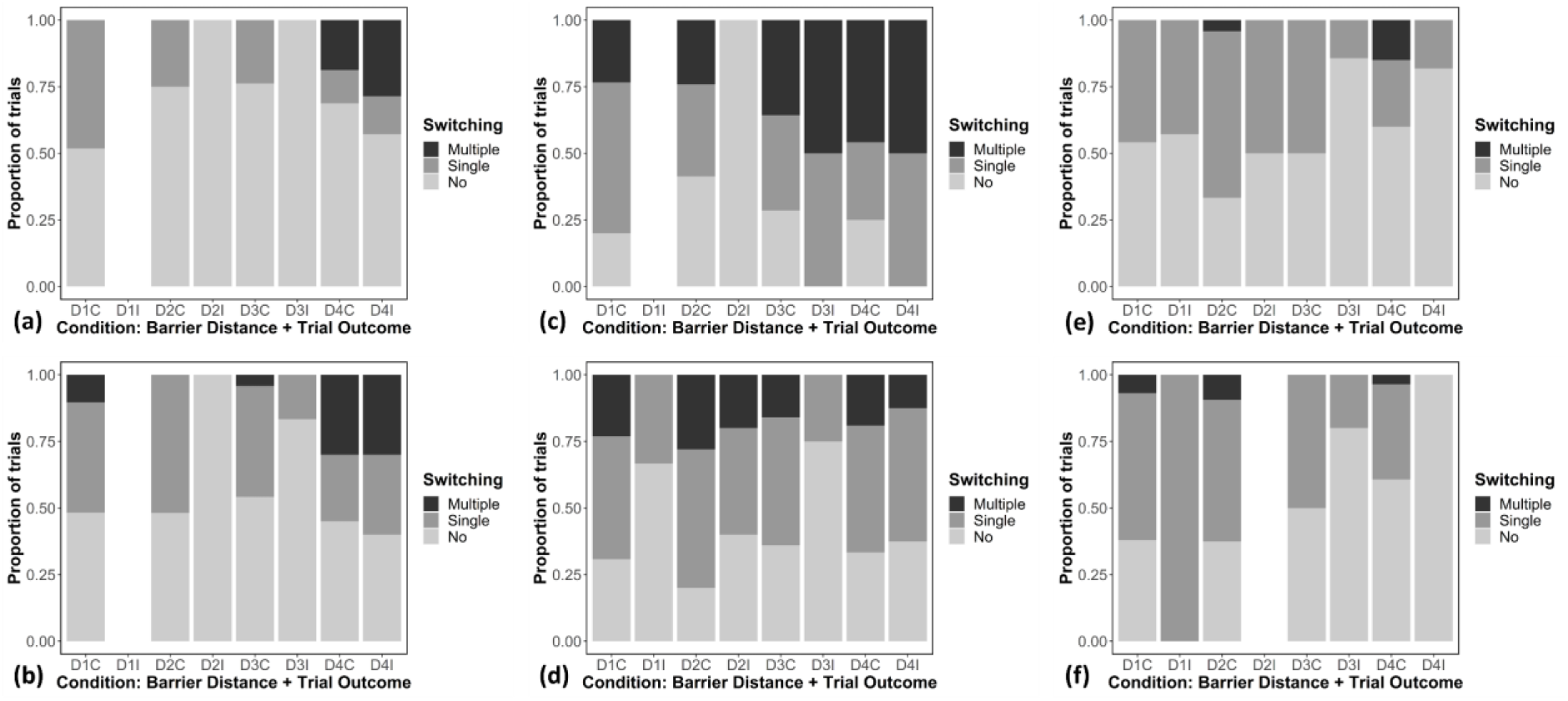
Side switching behaviour of fish one to six (a-f) during test probe trials. Trials have been separated according to barrier distance being tested (D1, D2, D3, or D4) and trial outcome (C=correct or I=correct). Greyscale bars indicate the type of side switching behaviour observed (lightest grey=no side switching, mid-grey=single side switch, and lastly, black=multiple (two or more) side switches). The size of each bar represents the proportion of trials which fitted each condition. Blank spaces denote conditions where no trials were available to assess. Note that fish five and six (e and f) were trained and tested without the doors by A and B. N trials: fish one=118 (a), fish two=118 (b), fish three=120 (c), fish four=117 (d), fish five=123 (e), and fish six=124 (f).

## References

Animal Behaviour Journal. (2018). Guidelines for the Use of Animals, Guidelines for the treatment of animals in behavioural research and teaching. Anim. Behav. 135, I–X.

Bates, D., Mächler, M., Bolker, B. M., & Walker, S. C. (2015). Fitting Linear Mixed-Effects Models Using lme4. J. Stat. Softw. 67: doi: 10.18637/jss.v067.i01

Caputi, A. A. (2004). Contributions of electric fish to the understanding sensory processing by reafferent systems. J. Physiol. Paris. 98. 81–97.

Caputi, A. A., & Budelli, R. (2006). Peripheral electrosensory imaging by weakly electric fish. J. Comp. Physiol. A Neuroethol. Sens. Neural. Behav. Physiol. 192, 587–600.

Engelmann, J., Bacelo, J., Metzen, M., Pusch, R., Bouton, B., Migliaro, A., … von der Emde, G. (2008). Electric imaging through active electrolocation: implications for the analysis of complex scenes. Biol. Cybern. 98, 519–539.

Engelmann, J., Wallach, A., & Maler, L. (2021). Linking active sensing and spatial learning in weakly electric fish. Curr. Opin. Neurobiol. 71, 1–10.

Fechler, K., & von der Emde, G. (2013). Figure–ground separation during active electrolocation in the weakly electric fish, Gnathonemus petersii. J. Physiol. Paris. 107, 72–83.

Francke, M., Kreysing, M., Mack, A., Engelmann, J., Karl, A., Makarov, F., … Reichenbach, A. (2014). Grouped retinae and tapetal cups in some Teleostian fish: Occurrence, structure, and function. Prog. Retin. Eye. Res. 38, 43–69.

Fujita, K., & Kashimori, Y. (2019). Representation of object’s shape by mutliple electric images in electrolocation. Biol. Cybern. 113, 239–255.

Gottwald, M., Bott, R. A., & von der Emde, G. (2017). Estimation of distance and electric impedance of capacitive objects in the weakly electric fish Gnathonemus petersii. J. Exp. Biol. 220, 3142–3153 doi:10.1242/jeb.159244

Gottwald, M., Sing, N., Haubrich, A. N., Regett, S., & von der Emde, G. (2018). Electric-color sensing in weakly electric fish suggests color perception as a sensory concept beyond vision. Curr. Biol. 28, 3648–3653, https://doi.org/10.1016/j.cub.2018.09.036

Heiligenberg, W. (1973). Electrolocation of Objects in the Electric Fish Eigenmannia (Rhamphichthyidae, Gymnotoidei). J. Comp. Physiol. 87, 137–164.

Hofmann, V., Sanguinetti-Scheck, J. I., Gómez-Sena, L., & Engelmann, J. (2013a). From static electric images to electric flow: Towards dynamic perceptual cues in active electroreception. J. Physiol. Paris. 107, 95–106.

Hofmann, V., Sanguinetti-Scheck, J. I., Künzel, S., Geurten, B., Gómez-Sena, L., & Engelmann, J. (2013b). Sensory flow shaped by active sensing: sensorimotor strategies in electric fish. J. Exp. Biol. 216, 2487–2500.

Hofmann, V., Sanguinetti-Scheck, J. I., Gómez-Sena, L., & Engelmann, J. (2017). Sensory Flow as a Basis for a Novel Distance Cue in Freely Behaving Electric Fish. J. Neurosci. 37(2), 302–312.

Kreysing, M., Pusch, R., Haverkate, D., Landsberger, M., Engelmann, J., Ruiter, J., … Francke, M. (2012). Photonic Crystal Light Collectors in Fish Retina Improve Vision in Turbid Water. Science. 336, 1700–1703.

Land, M. F. (1999). Motion and vision: why animals move their eyes. J. Comp. Physiol. A Neuroethol. Sens. Neural. Behav. Physiol. 185, 341–352. DOI: https://doi.org/10.1007/s003590050393, PMID: 10555268

Landsberger, M., von der Emde, G., Haverkate, D., Schuster, S., Gentsch, J., Ulbricht, E., … Wagner, H-J. (2008). Dim light vision - Morphological and functional adaptations of the eye of the mormyrid fish, Gnathonemus petersii. J. Physiol. Paris. 102, 291–303.

Lissmann, H. W. (1951). Continuous Electrical Signals from the Tail of a Fish, Gymnarchus niloticus Cuv. Nature. 167, 201–202.

Lissmann, H. W. (1958). On the function and evolution of electric organs in fish. J. Exp. Biol. 35, 156–191.

Lissmann, H. W., & Machin, K. E. (1958). The mechanism of object location in Gymnarchus niloticus and similar fish. J. Exp. Biol. 35, 451–486.

Mitchinson, B., Grant, R. A., Arkley, K., Rankov, V., Perkon, I., & Prescott, T. J. (2011). Active vibrissal sensing in rodents and marsupials. Phil. Trans. R. Soc. B. 366, 3037–3048.

Nelson, M. E., & MacIver, M. A. (2006). Sensory acquisition in active sensing systems. J. Comp. Physiol. A Neuroethol. Sens. Neural. Behav. Physiol. 192, 573–586.

Prescott, T. J., Diamond, M. E., & Wing, A. M. (2011). Active touch sensing. Phil. Trans. R. Soc. B. 366, 2989–2995.

Pusch, R., von der Emde, G., Hollmann, M., Bacelo, J., Nöbel, S., Grant, K., & Engelmann, J. (2008). Active sensing in a mormyrid fish: electric images and peripheral modifications of the signal carrier give evidence of dual foveation. J. Exp. Biol. 211, 921–934.

Pusch, R., Kassing, V., Riemer, U., Wagner, H-J., von der Emde, G., & Engelmann, J. (2013a). A grouped retina provides high temporal resolution in the weakly electric fish Gnathonemus petersii. J. Physiol. Paris. 107, 84–94.

Pusch, R., Wagner, H-J., von der Emde, G., & Engelmann, J. (2013b). Spatial Resolution of an Eye Containing a Grouped Retina: Ganglion Cell Morphology and Tectal Physiology in the Weakly Electric Fish Gnathonemus petersii. J. Comp. Neurol. 521. 4075–4093.

Schumacher, S., Burt de Perera, T., Thenert, J., & von der Emde, G. (2016a). Cross-modal object recognition and dynamic weighting of sensory inputs in a fish. Proc. Natl. Acad. Sci. U.S.A. 113(27), 7638–7643.

Schumacher, S., Burt de Perera, T., & von der Emde, G. (2016b). Object discrimination through active electrolocation: Shape recognition and the influence of electrical noise. J. Physiol.Paris. 110(3), 151–163.

Schumacher, S., Burt de Perera, T., & von der Emde, G. (2017a). Electrosensory capture during multisensory discrimination of nearby objects in the weakly electric fish Gnathonemus petersii. Sci. Rep. 7, 43665, doi: 10.1038/srep436651.

Schumacher, S., von der Emde, G., & Burt de Perera, T. (2017b). Sensory influence on navigation in the weakly electric fish Gnathonemus petersii. Anim. Behav. 132, 1–12.

Skeels, S. (2022). Sensory-motor integration in Gnathonemus petersii, PhD thesis. Oxford: University of Oxford.

Toerring, M. J., & Belbenoit, P. (1979). Motor programmes and electroreception in mormyrid fish. Behav. Ecol. Sociobiol. 4, 369–379.

Toerring, M-J., & Moller, P. (1984). Locomotor and electric displays associated with electrolocation during exploratory behavior in mormyrid fish. Behav. Brain Res. 12, 291–306.

von der Emde, G., Schwarz, S., Gomez, L., Budelli, R., & Grant, K. (1998). Electric fish measure distance in the dark. Nature. 395, 890–894.

von der Emde, G., & Schwarz, S. (2000). Three–dimensional analysis of object properties during active electrolocation in mormyrid weakly electric fishes (Gnathonemus petersii). Philos. Trans. R. Soc. B. 335(1401), 1143–1146.

von der Emde, G., & Schwarz, S. (2002). Imaging of Objects through active electrolocation in Gnathonemus petersii. J. Physiol. Paris. 96, 431–444.

von der Emde, G. (2004). Distance and shape: perception of the 3-dimensional world by weakly electric fish. J. Physiol. Paris. 98, 67–80.

von der Emde, G. (2006). Non-visual environmental imaging and object detection through active electrolocation in weakly electric fish. J. Comp. Physiol. A Neuroethol. Sens. Neural. Behav. Physiol. 192, 601–612.

von der Emde, G., & Fetz, S. (2007). Distance, shape and more: Recognition of object features during active electrolocation in a weakly electric fish. J. Exp. Biol. 210(17), 3082–3095.

von der Emde, G., Behr, K., Bouton, B., Engelmann, J., Fetz, S., & Folde, C. (2010). 3-Dimensional scene perception during active electrolocation in a weakly electric pulse fish. Front. Behav. Neurosci. 4, Article 26, doi: 10.3389/fnbeh.2010.00026.

von der Emde, G., & Zeymer, M. (2020). Multisensory object detection in weakly electric fish. In The Senses: A Comprehensive Reference (2nd edition): 1–17. Fritsch, B. (Ed.). Elsevier Inc. https://doi.org/10.1016/B978-0-12-809324-5.24211-9

Yang, S. C-H., Wolpert, D. M., & Lengyel, M. (2016). Theoretical perspectives on active sensing. Curr. Opin. Behav. Sci. 11, 100–108.

